# *Effect of host-mimicking medium and biofilm growth on the ability of colistin to kill* Pseudomonas aeruginosa

**DOI:** 10.1101/2020.08.25.265942

**Authors:** Esther Sweeney, Akshay Sabnis, Andrew M. Edwards, Freya Harrison

## Abstract

*In vivo* biofilms cause recalcitrant infections with extensive and unpredictable antibiotic tolerance. Here, we demonstrate increased tolerance of colistin by *Pseudomonas aeruginosa* when grown in cystic fibrosis-mimicking medium *versus* standard medium in *in vitro* biofilm assays, and drastically increased tolerance when grown in an *ex vivo* CF model versus the *in vitro* assay. We used colistin conjugated to the fluorescent dye BODIPY to assess the penetration of the antibiotic into *ex vivo* biofilms and showed that poor penetration partly explains the high doses of drug necessary to kill bacteria in these biofilms. The ability of antibiotics to penetrate the biofilm matrix is key to their clinical success, but hard to measure. Our results demonstrate both the importance of reduced entry into the matrix in *in vivo*-like biofilm, and the tractability of using a fluorescent tag and benchtop fluorimeter to assess antibiotic entry into biofilms. This method could be a relatively quick, cheap and useful addition to diagnostic and R&D pipelines, allowing the assessment of drug entry into biofilms, in *in vivo*-like conditions, prior to more detailed tests of biofilm killing.

## Full text

Biofilm infections of host tissues or indwelling medical devices impose a significant health and economic burden, due to the high tolerance of biofilm bacteria to host immune attack and to antibiotics. Biofilm antibiotic tolerance is a function of environmentally-cued changes in bacterial physiology and gene expression, and reduced penetration of some antibiotic molecules through the biofilm matrix (1). In the case of cystic fibrosis (CF) lung disease, plugging of small airways by aggregates of biofilm embedded in abnormal host mucus leads to reduced airflow and bronchiectasis (2, 3). In a recent analysis, bacterial lung infection was the strongest predictor of medication costs in CF, adding on average €3.6K per patient per year to direct healthcare costs (4, 5). Intravenous antibiotic treatments, usually administered during acute exacerbations of respiratory symptoms, impose a particularly heavy burden: in the UK, people with CF spend a median of 27 days/year receiving IV antibiotics, often as hospital in-patients (6). There is a narrow choice of antibiotics suitable for administration in CF, and a poor concordance between antibiotic susceptibility testing in diagnostic labs and patient outcome – even when standard *in vitro* biofilm platforms are employed for testing (7, 8).

Generic *in vitro* biofilm models (e.g. Calgary device (9, 10), flow cells) and standard laboratory growth media are usually used to test the efficacy of antibacterial agents, both in a diagnostic setting and in R&D pipelines for new agents. It is increasingly recognised that the key to optimising biofilm management is a more context-specific approach to mimicking the in-host conditions in the infection(s) of interest. The effect of physicochemical environment, and especially of host-mimicking *versus* standard laboratory media, on bacterial responses to antibiotics is well known (11-14). Further, environmental differences between specific biofilm contexts are likely to produce different biofilm architectures; overall biofilm thickness and 3D structure, as well as the production of matrix polymers with different size, charge or hydrophobicity, will impact how easily different antibiotics can penetrate the matrix to reach cells (15). The benefits and shortcomings of different *in vitro* and *in vivo* biofilm models, and limitations on how well standard models such as the Calgary device represent *in vivo* biofilms, have been reviewed in detail by other authors (16-18).

We hypothesised that poor penetration of the *in vivo* matrix partly explains the high doses of antibiotic necessary to kill *Pseudomonas aeruginosa* in CF lung biofilms. We focussed on the antibiotic colistin for two reasons. First, colistin is widely prescribed for *P. aeruginosa* infection in CF and is commonly administered by inhalation; this means that topical lab exposure of biofilms to colistin, by spiking the surrounding culture medium, likely mimics *in vivo* exposure better than in the case of antibiotics that are only administered orally or through IV. Second, fluorescently-labelled colistin was available from colleagues, opening up the possibility to assay colistin concentration in biofilm via fluorimetry (19). Finally, colistin is able to bind the *P. aeruginosa* exopolymer Psl (20) and extracellular lipopolysaccharide (21), and may adsorb to outer membrane vesicles (22) – all of which may be present in *P. aeruginosa* biofilms depending on strain and culture conditions. We aimed first to compare the tolerance of *P. aeruginosa* to colistin using planktonic microdilution in standard antibiotic susceptibility testing medium (cation-adjusted Muller-Hinton broth, caMHB) and synthetic CF sputum medium (SCFM, (23)); biofilm eradication assays using the same media in a Calgary device; and in an *ex vivo* model of CF biofilm which combines SCFM and pig bronchiolar tissue (24, 25). We then aimed to use fluorescently-labelled colistin to measure the percentage of a dose which was able to enter the biofilm matrix in the *ex vivo* CF model.

The lab strains PAO1 (Nottingham isolate) and PA14 (a gift from Leo Eberl) and eight isolates of *P. aeruginosa* from a chronically-colonised person with CF (SED6, SED7, SED8, SED9, SED11, SED13, SED17, SED19 (26)) were used in this work. The CF strains had previously been shown to vary considerably in antibiotic resistance profile, and in biofilm forming ability as measured by crystal violet staining following growth in Lysogeny Broth (LB) in a Calgary biofilm device (S. Darch, *pers. commun*.) Colonies of frozen stocks were obtained by growth on LB agar at 37°C for 18-24h (PAO1, PA14) or 48h (CF isolates).

The minimum inhibitory concentration (MIC) and minimum biofilm eradication concentration (MBEC) of colistin (Acros organics) for the strains was assayed by Warwick’s Antimicrobial Screening Facility following CLSI guidelines, in cation-adjusted Müller-Hinton broth and in SCFM. SCFM was made following the recipe of Palmer *et al*. (23), with the modification that glucose was removed because previous work suggested that this promoted the growth of endogenous bacteria present on pig lung tissue. MIC testing was performed according to CLSI guidelines M7-A9 (Methods for Bacteria that Grow Aerobically), M24-A (Testing of Mycobacteria, Nocardiae, and Other Aerobic Actinomycetes), and M100-S24 (Performance Standards for Antimicrobial Susceptibility Testing). MBEC testing used the Calgary Biofilm Device (peg lid assay) using methods published by Moskowitz *et al*. (10). As expected, the inhibitory concentration of colistin varied depending on culture medium (SCFM usually > caMHB) and growth mode (MBEC > MIC in all cases but one), Figure 1. For 6 of the 9 isolates, conducting MIC testing in SCFM instead of caMHB led to a change in classification from sensitive to resistant. We selected CF isolates SED6, SED8, SED17 and SED19 for further work with the *ex vivo* pig lung model, as these had a range of MIC/MBEC values and included two isolates for which MBEC=MIC in SCFM (SED6, SED8) and two for which MBEC>MIC in SCFM (SED17, SED19).

**Figure 1.**
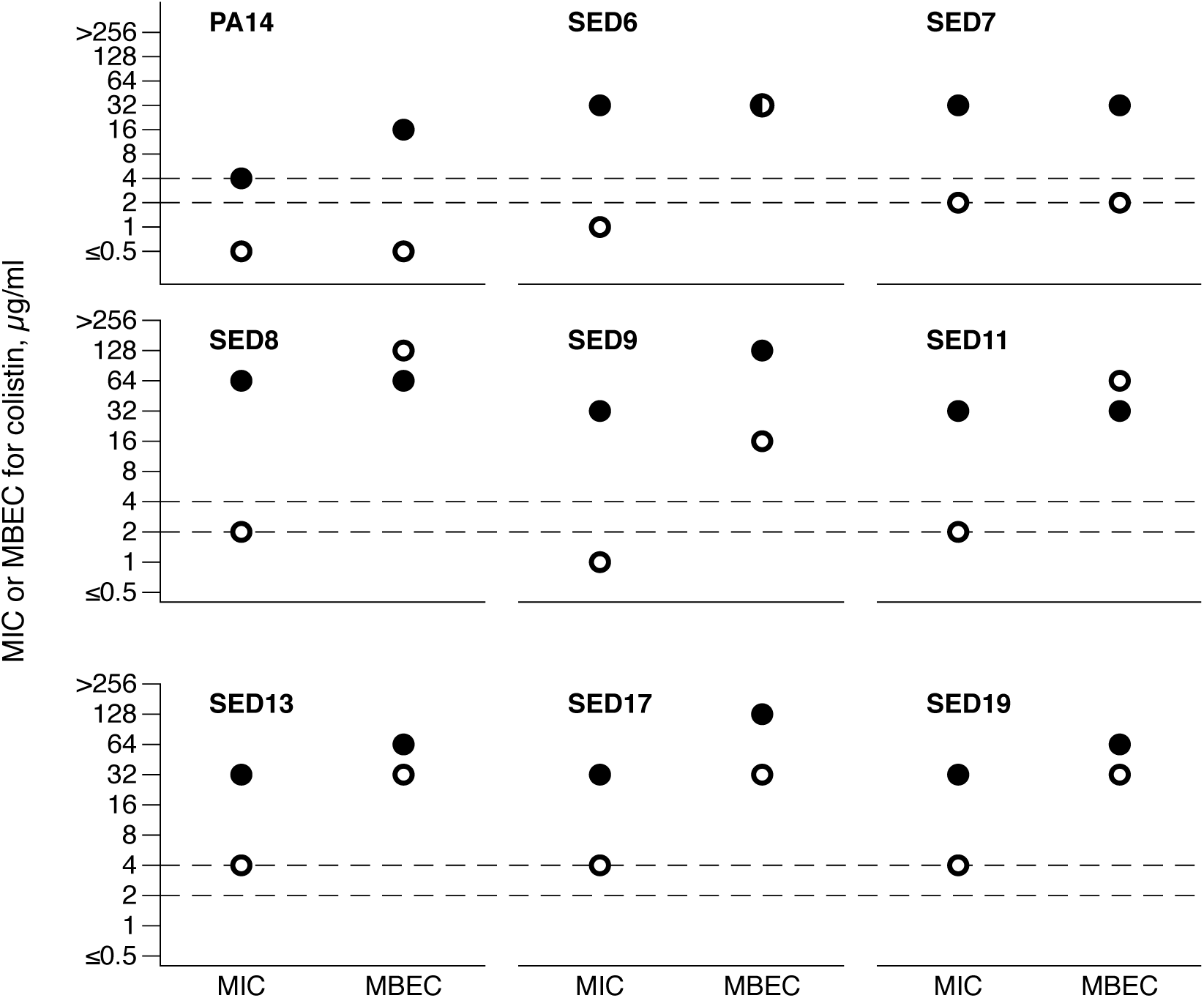
MIC and MBEC of colistin for PA14 and eight CF isolates of *P. aeruginosa*. Open symbols show values for tests conducted in cation-adjusted Müller-Hinton broth, closed symbols show values for tests conducted in synthetic CF sputum medium; note that for SED 8, the MBEC was the same in both media. The dashed lines denote the CLSI breakpoints for classification as sensitive (≤2 µg.ml^-1^) or resistant (≥ 4 µg.ml^-1^) in MIC testing in cation-adjusted Müller-Hinton broth. Full data are supplied in the Data Supplement; MIC values are taken from two replica assays and MBECs are from a single assay.

The *ex vivo* pig lung model (EVPL) was prepared as previously described (24, 25), except that SCFM was not supplemented with antibiotics to repress the growth of any endogenous bacteria present on the lung tissue: this was to prevent any interference with assessment of colistin efficacy in later experiments. Full details are in the supplementary methods. *P. aeruginosa* has been shown to produce CF-like biofilm in this model (24).

Reproducibility of biofilm load is essential in any model system which is intended for use in testing of antibiofilm or antibacterial agents. We therefore assessed the reproducibility of biofilm load between pieces of tissue from same/different lungs, by measuring the colony-forming units (CFU, by plating on LB agar) in biofilms of PAO1 and PA14 at 2 and 7 days post infection. Figure S1 shows photographs of tissues at inoculation and at 2 and 7 days p.i.; *P. aeruginosa* growth is clearly visible due to the typical blue-green pigmentation of this species. All colonies recovered from tissue infected with *P. aeruginosa* were identified as *P. aeruginosa* by morphology; uninfected tissues cultured a high load of endogenous bacteria from the lungs but these were clearly distinguishable from *P. aeruginosa* by size, shape and colour. As shown in Figure S2, both PAO1 and PA14 reached biofilm loads comparable with those reported for cystic fibrosis (typical figures in the literature are around 10^8^ CFU per ml of sputum). As detailed in Figure S3, Table S1 and the supplementary methods, PA14 showed more reproducible biofilm loads than PAO1; biofilm loads were more repeatable at day 2 than day 7. Standardising CFU by tissue area (measured in mm^2^ using ImageJ, (27)) did not improve repeatability, given the variation inherent in cutting tissue by hand (see Supplementary Data file) this result was initially surprising. However, the tissue may simply be a scaffold, or cue, for physiologically realistic biofilm formation & maturation, rather than a nutrient source. If bacteria gain most of their nutrients from the ASM rather than the tissue, attempting to standardise by tissue area is unnecessary – the bacteria have reached their carrying capacity so dividing this by area, which varies considerably, adds stochastic variation. Based on these preliminary results we decided to use PA14 as an example lab isolate in further work; to grow biofilms for two days prior to colistin treatment; and not to standardise CFU counts by tissue area when measuring viable cells in biofilms.

## Abbreviations

caMHB: Muller-Hinton broth
CF: cystic fibrosis
CLSI: Clinical and Laboratory Standards Institute
EUCAST: European Committee on Antimicrobial Susceptibility Testing
EVPL: *ex vivo* pig lung
LB: lysogeny broth
MBC: minimum bactericidal concentration
MIC: minimum inhibitory concentration
MBEC: minimum biofilm eradication concentration
SCFM: synthetic cystic fibrosis sputum medium

## Supplementary methods

### Preparation of EVPL

Pig lungs were obtained from Quigley & Sons, Cubbington and John Taylor & Son, Earlsdon and dissected on the day of delivery under sterile conditions. The pleura of the ventral surface was heat sterilised using a hot pallet knife. A sterile razor blade was then used to make an incision in the lung, exposing the bronchiole. A section of the bronchiole was extracted and the exterior alveolar tissue removed using dissection scissors. Bronchiolar sections were washed in a 1:1 mix of Dulbecco’s modified Eagle medium (DMEM) and RPMI 1640 supplemented with 50 µg ml^-1^ ampicillin (Sigma-Aldrich). Bronchioles were then cut into squares, with a further 2 washes in DMEM+RPMI+ampicillin during dissection. The bronchiole squares were then washed in SCFM (no antibiotics), UV sterilised for 5 minutes in a germicidal cabinet and transferred to individual wells of a 24-well plate containing 400 µl

SCFM solidified with 0.8% (w/v) agarose per well. Tissue sections were inoculated by touching a sterile 29G hypodermic needle (Becton Dickinson Medical) to the surface of a *P. aeruginosa* colony on LB agar, and gently piercing the surface of the tissue section. Uninfected control sections were mock inoculated with a sterile needle. 500 µl SCFM was added to each well. Unlike our initial published work on this model, we did not supplement SCFM with antibiotics to repress the growth of any endogenous bacteria present on the lung tissue. This was because we did not want this to interfere with assessment of colistin efficacy in later experiments. Plates were sealed with Breathe-Easier^®^ gas-permeable membrane (Diversified Biotech) and incubated at 37°C for the desired length of time.

### Assessing repeatability of *P. aeruginosa* biofilm load on *ex vivo* pig bronchiole

Eighteen bronchiolar sections from each of 3 sets of pig lungs were prepared as above. 24-well plates containing tissues were photographed using a 20MP digital camera placed at a set height on a copy stand and these images later used to calculate the surface area of each tissue section using the freehand select tool in ImageJ: as we dissect tissues by hand we wanted to determine whether variation in section size affected bacterial load. The mean size of tissue section was 44 mm^2^ (s.d. 11.5 mm^2^). Six sections of tissue were inoculated with PA14 and six with PAO1; the remaining six were left uninoculated. After 2 and 7 days incubation at 37°C, half of the tissue sections plus associated biofilm were removed, briefly washed in 500 µl PBS in a fresh 24-well plate to remove loosely-adhering planktonic cells, then placed into 1ml PBS in screw-cap homogenisation tubes (Fisherbrand) containing eighteen 2.38 mm metal beads (Fisherbrand). Tissue was bead beaten in a FastPrep-24 5G (MP Biomedicals) for 40 s at 4 m s^-1^ to recover the bacteria from the tissue-associated biofilm. Bacterial load was calculated by serially diluting the homogenates, plating on LB agar and counting colony-forming units (CFU).

We wished first to address two questions. First, whether there was a statistically significant interaction between lung and *P. aeruginosa* strain in determining total CFU or CFU/mm^2^, i.e. did any difference between PA14 and PAO1 depend on whether they were in tissue from lung 1, 2 or 3? A lack of such an interaction would suggest that experimental results are robust to any natural variation in tissue between lungs taken from different animals. Second, we wished to calculate the statistical repeatability of total CFU or CFU/mm^2^ across lungs 1-3 for each of the two strains as a measure of reproducibility. Based on the answers to these two questions, we then aimed to make decision about (1) whether future experiments should measure the effects of antibiotics in terms of total CFU or whether we should standardise for variation in the size of hand-cut tissue sections by working with counts of CFU/mm^2^; and (2) whether one of the two lab strains grew more consistently in the model and thus would be better to use as a standard “wild type” in future work.

### Production of labelled colistin

This process is described in detail by Sabnis *et al*. (19). Briefly, colistin was labelled by incubating with BODIPY® FL SE D2184 (Thermo Fisher Scientific) and sodium bicarbonate for 2 h at 37°C; unbound BODIPY was removed by dialysis; and successful labelling was confirmed by time-of-flight mass spectrometry.

**Figure S1.**
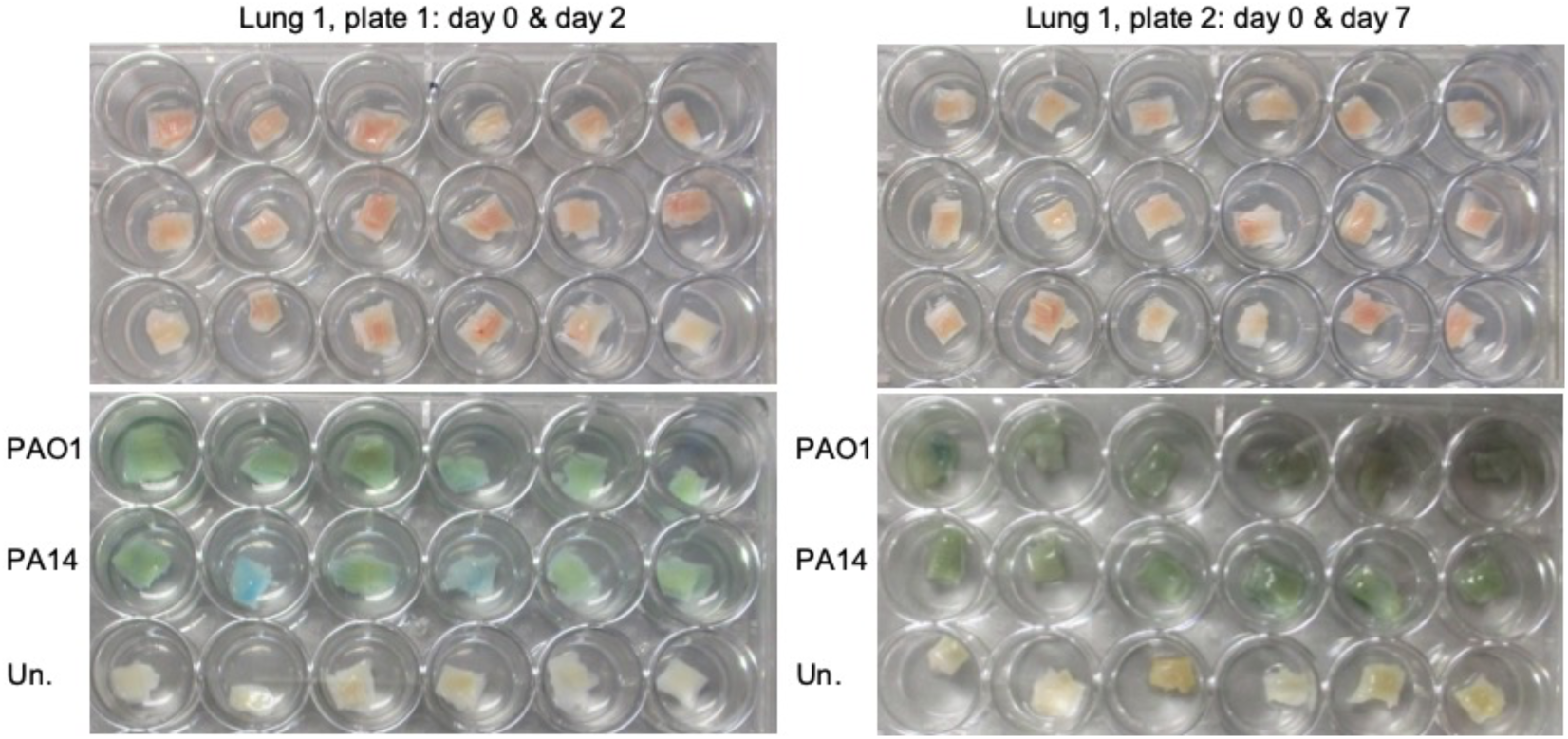
Photographs of example tissue sections and biofilms. Sections of pig bronchiole from one lung are shown in standard 24-well culture plates prior to inoculation (top panels) and after either two or seven days ‘incubation at 37°C.

**Figure S2.**
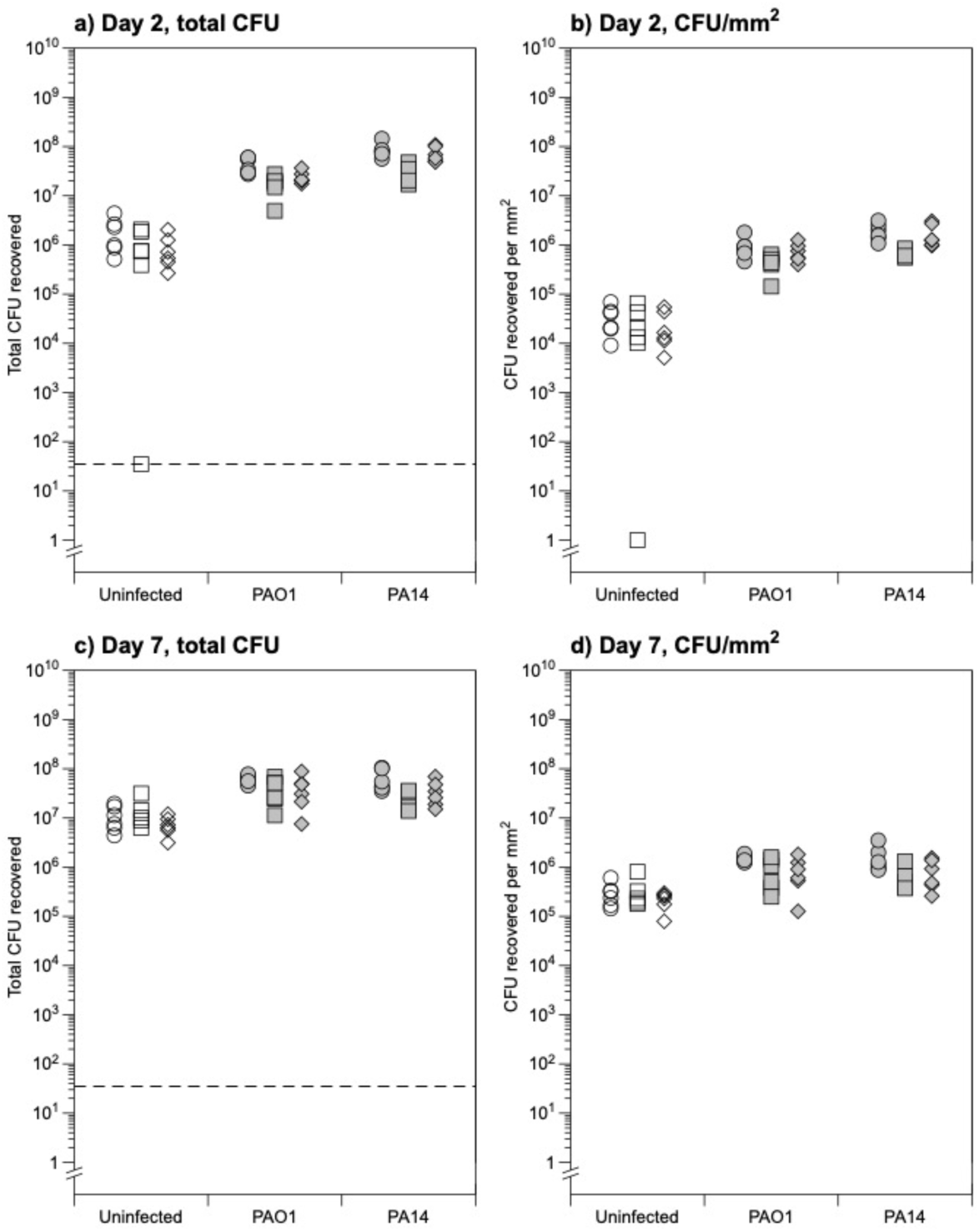
Bacterial CFU counts in biofilm harvested from tissue sections at 2 and 7 days post infection. Open symbols denote colonies of endogenous bacteria, closed symbols colonies identifiable as *P. aeruginosa*. Circles: lungs 1; squares: lung 2, diamonds: lung 3. ANOVA showed there was no significant interaction between lung and strain at either day, in total or area-standardised data (see Table S1). Raw data, R code and full results of ANOVA analyses of these data are supplied in the Supplementary Data file.

**Table S1.**
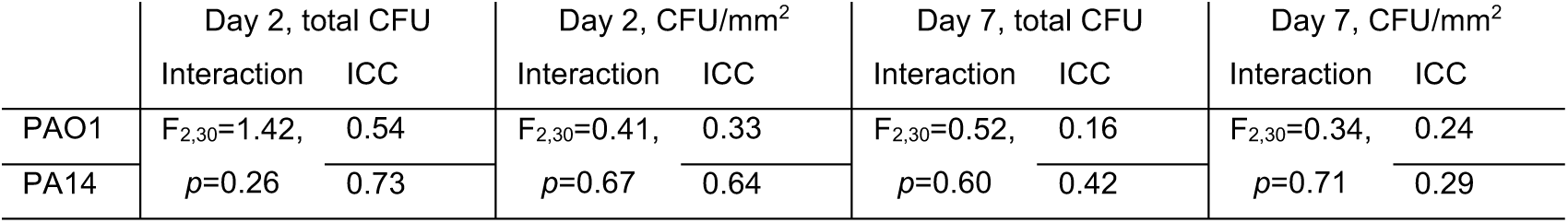
Reproducibility of biofilm loads on *ex vivo* tissue. The data in Figure S2 were analysed using ANOVA to test for effects of lung, strain and their interaction on bacterial load, and by linear mixed-effects models to calculate the variance in each species ‘bacterial load within and between lungs. This latter information was used to calculate the intraclass correlation coefficient – this is simply the proportion of total variance explained by lung and is a commonly-used measure of repeatability. The larger this value, the greater the between-lung variance relative to the within-lung variance, and so the less noise present in the data due to random variation in biofilm load between tissue sections (“error”). In the table, “Interaction” records results for the lung*strain interaction term in ANOVAs. “ICC” refers to the intra-class correlation coefficient. Raw data, R code and full results of analyses are supplied in the Supplementary Data file.

We then aimed to assess the bactericidal activity of colistin to EVPL-grown *P. aeruginosa*, and identify concentrations that were sub-inhibitory or bactericidal to these biofilms for use in later experiments with labelled colistin. PA14 and clinical isolates SED6, SED8, SED17 and SED19 were cultured in EVPL+SCFM for 2 days, and tissue pieces plus surrounding biofilm then transferred to new 24-well plates containing 1 ml fresh SCFM+colistin at 0.5MIC, 4xMIC, 0.5xMBEC, 4xMBEC and 10xMBEC, or antibiotic-free SCFM (three pieces of tissue per strain, per treatment). The tissue sections + biofilm were incubated in these treatment plates for 18h at 37°C, and then bacteria were recovered, bead beaten as above and diluted and plated on LB agar for enumeration of viable CFU. The results in Figure 2 show that growth as biofilms in EVPL leads to a drastic increase in colistin tolerance compared with growth in *in vitro* surface-attached biofilm models, even when SCFM was used in conjunction with the *in vitro* MBEC platform. This is potentially due to decreased bacterial sensitivity, low penetration of colistin into the biofilm matrix and/or colistin binding proteins in the bronchiolar tissue (28)

**Figure 2.**
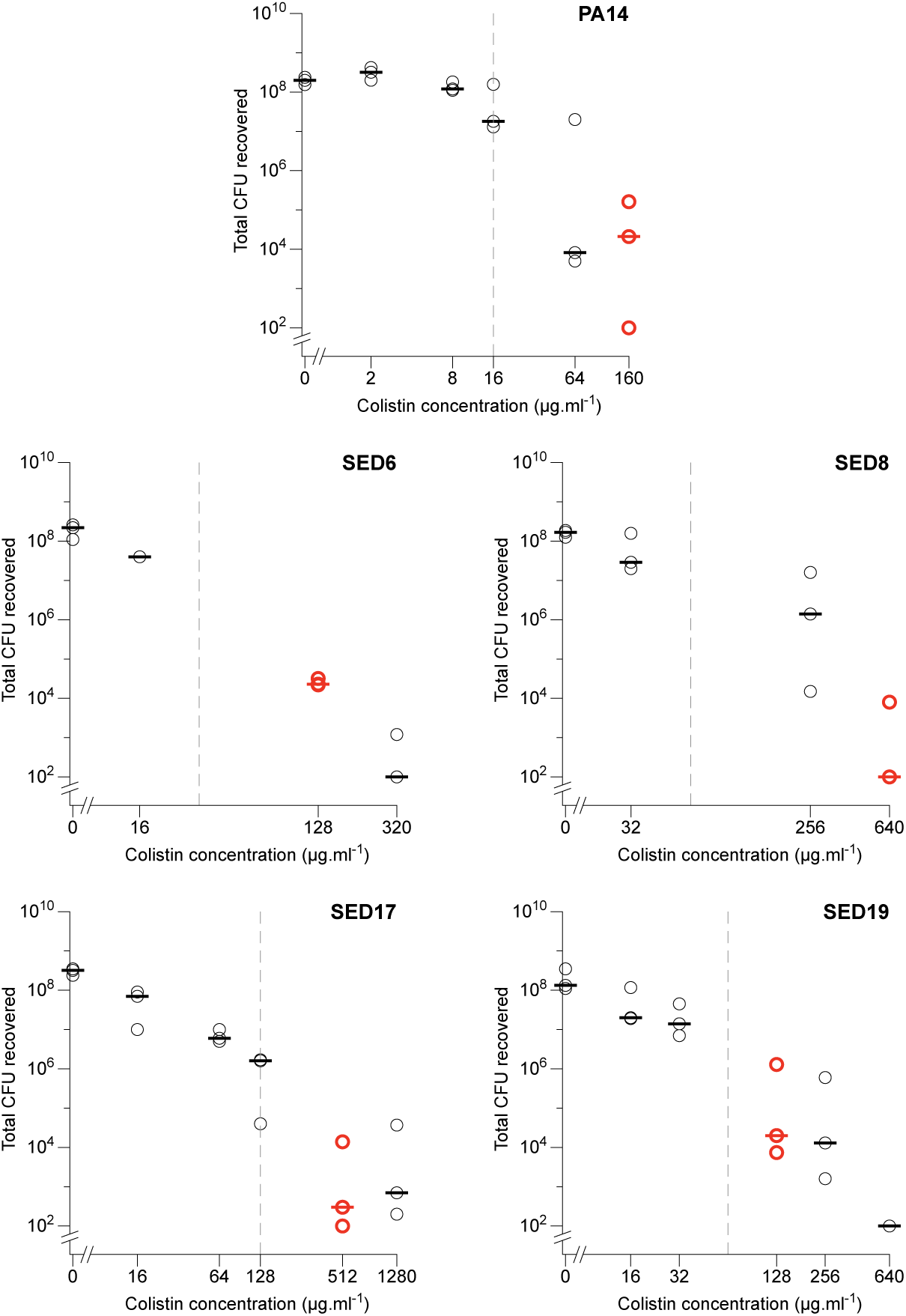
Effect of treating two-day EVPL biofilms of *P. aeruginosa* with colistin for 18h. Each data point represents a single tissue section, and all tissue sections were taken from the same pair of pig lungs. Horizontal lines denote medians. Red points show the lowest concentration of colistin causing a ≥3-log_10_ reduction in CFU compared with untreated biofilms (an outlier for PA14 at 64µg.ml^-1^ was discounted). PA14, SED17 and SED19 had differing values for MIC and MBEC tests in SCFM, and so were treated with colistin at 0.5MIC, 4xMIC, 0.5xMBEC, 4xMBEC and 10xMBEC. SED6 and SED8 had equal MIC and MBEC values in SCFM and so were treated with colistin at 0.5MIC/MBEC, 4xMIC/MBEC and 10xMIC/MBEC. Grey dashed lines show each isolates’s MBEC in SCFM in the Calgary device. Note that tissues + biofilms were exposed to BODIPY-colistin in a total volume of 1 ml SCFM, therefore concentrations correspond to total µg present. Raw data, R code and results of ANOVA analyses CFU in untreated biofilms formed by each strain are supplied in the Data Supplement.

Following CLSI recommendations (M26-A, Methods for Determining Bactericidal Activity of Antimicrobial Agents) the minimum bactericidal concentration (MBC) of colistin in the EVPL was defined as the lowest concentration causing a ≥3-log_10_ reduction in median CFU recovered from biofilms, compared with untreated biofilms (red circles in Figure 2). These concentrations were chosen as “bactericidal” concentrations of BODIPY-colistin to use in fluorescent colistin penetration assays (note that addition of BODIPY to colistin renders it inactive as an antibiotic). This corresponded to 10× the ASM MBEC for PA14 and SED8, 4× ASM MBEC for SED6 and SED17, and 2× SCFM MBEC for SED19. MBCs differed five-fold between the strains used. This is likely due to differences in biofilm matrix structure or other genetic differences between the isolates, as all grew to similar cell densities in the absence of colistin (ANOVA for effect of strain identity on *P. aeruginosa* CFU in untreated biofilms only: F_4,10_ = 1.49, *p* = 0.277). We chose 2µg ml^-1^ as the sub-inhibitory concentration for later work with PA14, and 8µg ml^-1^ as the sub-inhibitory concentration for later work with all CF isolates.

PA14, SED6, SED8, SED17 and SED19 were inoculated into replica pieces of EVPL, incubated for two days at 37°C to form mature biofilm, and tissue pieces then transferred individually to 30ml soda glass screw-top vial containing 1 ml fresh SCFM alone or 1 ml SCFM + colistin labelled with BODIPY FL SE D2184 (succinimidyl ester - Thermo Fisher Scientific) at concentrations corresponding to sub-inhibitory (2µg ml^-1^ for PA14, 8µg ml^-1^ for the CF isolates) or bactericidal concentrations (see Figure 2) of unlabelled colistin. Three uninfected pieces of EVPL were incubated alongside the infected tissue sections, and transferred individually to 30ml soda glass screw-top vial containing 1 ml fresh SCFM alone. Labelled colistin was a gift from Akshay Sabnis and Andrew Edwards (see (19)) Tubes were incubated for 18h at 37°C. Glass vials were used for these experiments because colistin can bind plastic and we wished to minimise loss to binding of the culture vessel in order to measure the proportion of colistin entering the biofilms as accurately as possible. Addition of the BODIPY-tag has been shown to reduce bactericidal activity in planktonic culture (19), and a pilot experiment using PA14, SED6 and SED8 had confirmed that the tag significantly reduced activity in our biofilm assay (Figure S3).

Tissue + biofilm was removed from the glass tubes and homogenised in 1 ml SCFM. Replica aliquots of this homogenate were used for CFU plating. BODIPY-colistin was immediately quantified in 100µl aliquots of tissue + biofilm homogenate and surrounding SCFM by fluorimetry (100µl in a black 96-well plate with ex/em 485/535 nm in a Tecan SPARK 10M multimode plate reader). There was slightly more background fluorescence signal from infected but untreated EVPL sections than from uninfected untreated EVPL (see Data Supplement), therefore fluorescence values from samples exposed to BODIPY-colistin were standardised by subtracting the mean fluorescence measured in homogenate/surrounding ASM from tissues infected by the same strain but exposed to SCFM with no BODIPY-colistin. A calibration curve using BODIPY-colistin in SCFM (Figure S5) was used to calculate total amounts of BODIPY-colistin in the whole biofilm and whole volume of surrounding SCFM. The percentage of the original dose of BODIPY-colistin recovered from both biofilm and SCFM combined was then calculated, to determine if any of the dose had been lost due to binding the glass, lung tissue and/or homogenisation beads.

**Figure S3.**
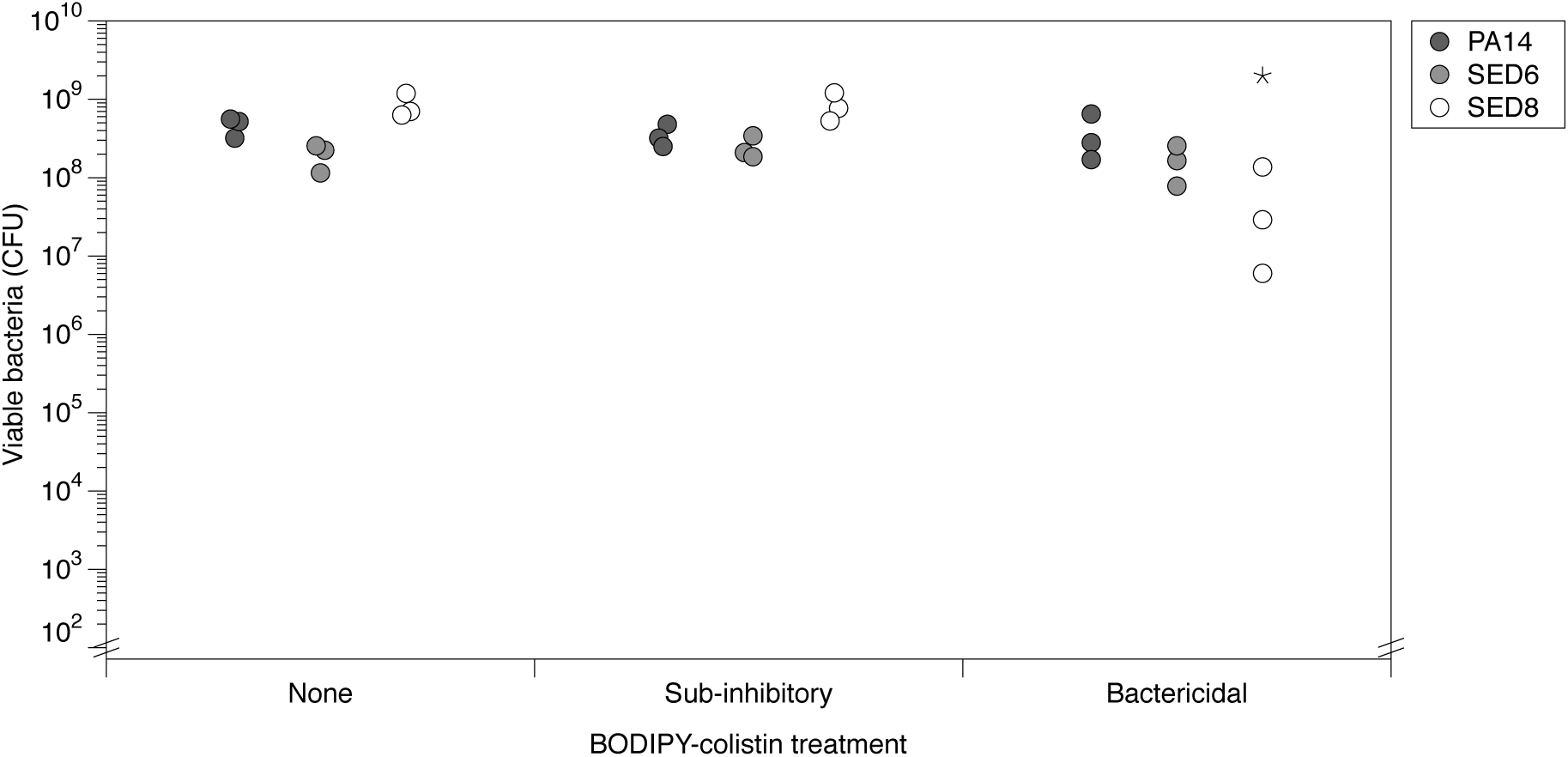
Effect of BODIPY-colistin on viable cells recovered from EVPL biofilms. SED6, PA14 and SED8 were selected as examples of strains with low, medium and high (128, 160 and 640 µg.ml^-1^) bactericidal concentrations of BODIPY-colistin, respectively. Aliquots of homogenised tissue+biofilm from these strains was diluted and plated for CFU counting after the exposure period. The graph shows CFU recovered from individual tissue sections. ANOVA was used to test for effects of strain, treatment (no, subinhibitory or bactericidal concentration) and their interaction. The residual mean square from the ANOVA was used to conduct planned pairwise t-tests to compare the mean CFU from biofilm exposed to BODIPY-colistin with the CFU from biofilms of the same strain that were not treated with BODIPY-colistin. A significant drop in CFU was observed only for SED8 treated with the highest concentration (denoted with *; t_18_ = 6.23, *p* < 0.001; CFU was approx. 20% of that observed in untreated biofilms), all other comparisons were not significant. Full data and statistical analysis are supplied in the Data Supplement.

**Figure S4.**
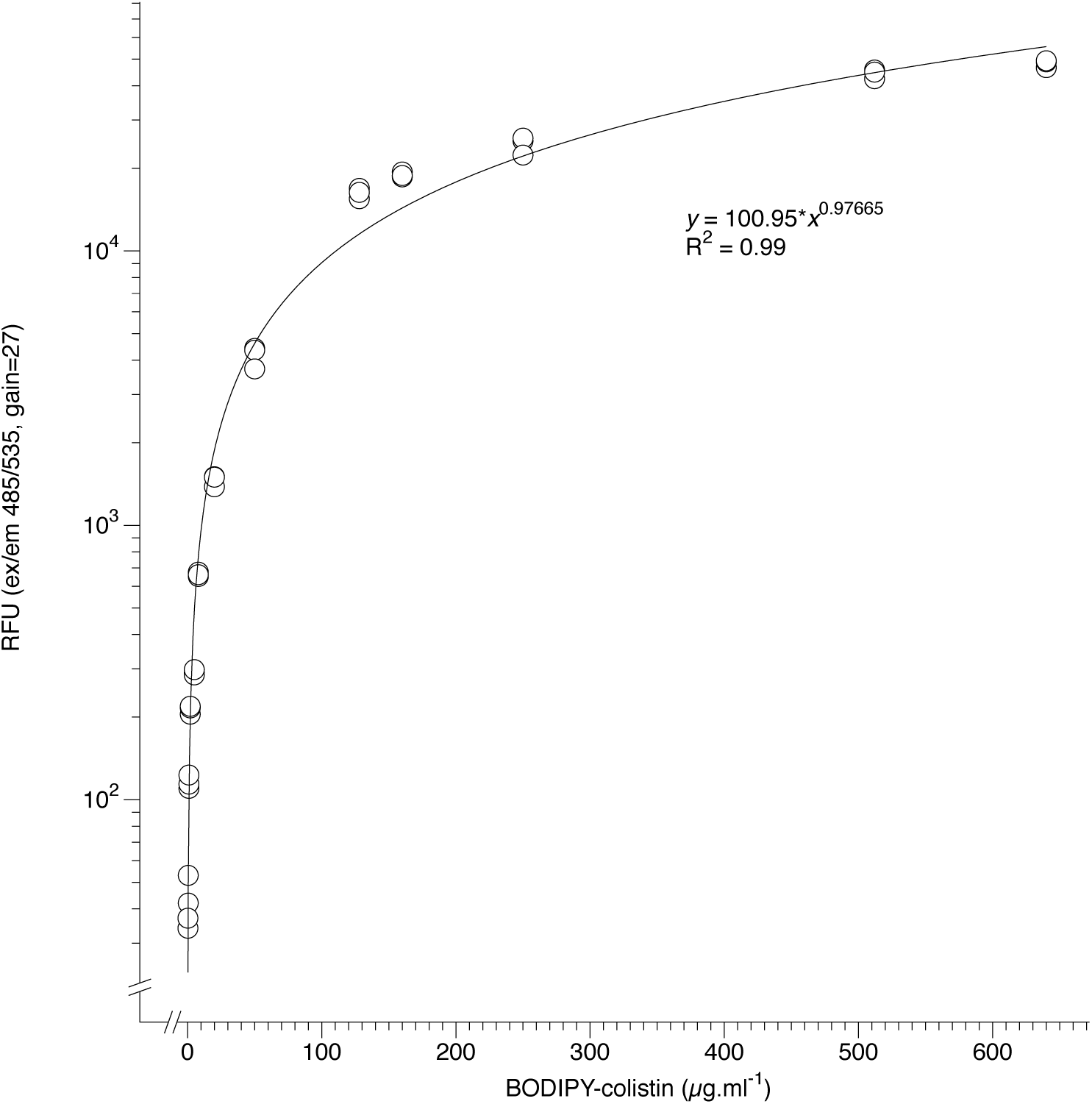
Calibration curve for BODIPY-colistin fluorescence. BODIPY-colistin was diluted in SCFM (0-640µg.ml^-1^) and incubated for 18h at 37°C before fluorescence (relative fluoresce units, RFU) was measured for triplicate 100µl aliquots, plus triplicate aliquots of SCFM only, in a Tecan SPARK 10M. The incubation period was the same as for the tissue sections exposed to BODIPY-colistin to allow for comparable degradation of the signal due to time or temperature in both experimental samples and calibration samples. The best fit was calculated in DataGraph 4.5.1 (Visual Data Tools Inc.) with equal weighting for all data points and R^2^ was 0.99. Raw data is supplied in the Data Supplement.

Recovery rates of total BODIPY-colistin from the biofilm homogenate plus surrounding SCFM were fairly good for medium-high concentrations, peaking at 93-100% recovery from lung sections receiving a dose of 128µg.ml^-1^ (Figure S5). At lower concentrations, recovery rates dropped, and became very variable at the lowest dose of 2 µg.ml^-1^ (used for PA14). This means that the values for this specific treatment may be unreliable. Our recovery rate of 70-80% at the second lowest dose (8 µg.ml^-1-^) was consistent with recovery rates in soda glass reported by a previous study which compared the loss of colistin to substrate binding in various materials (29). Better rates were achieved for higher concentrations, although recovery did then start to tail off as concentrations became very high. A pilot experiment in which the BODIPY-colistin exposure was conducted in plastic 24-well plates (Corning Costar®) resulted in only approx. 50% of the original BODIPY-colistin dose being recovered, consistent with colistin binding plastic (data not shown).

**Figure S5.**
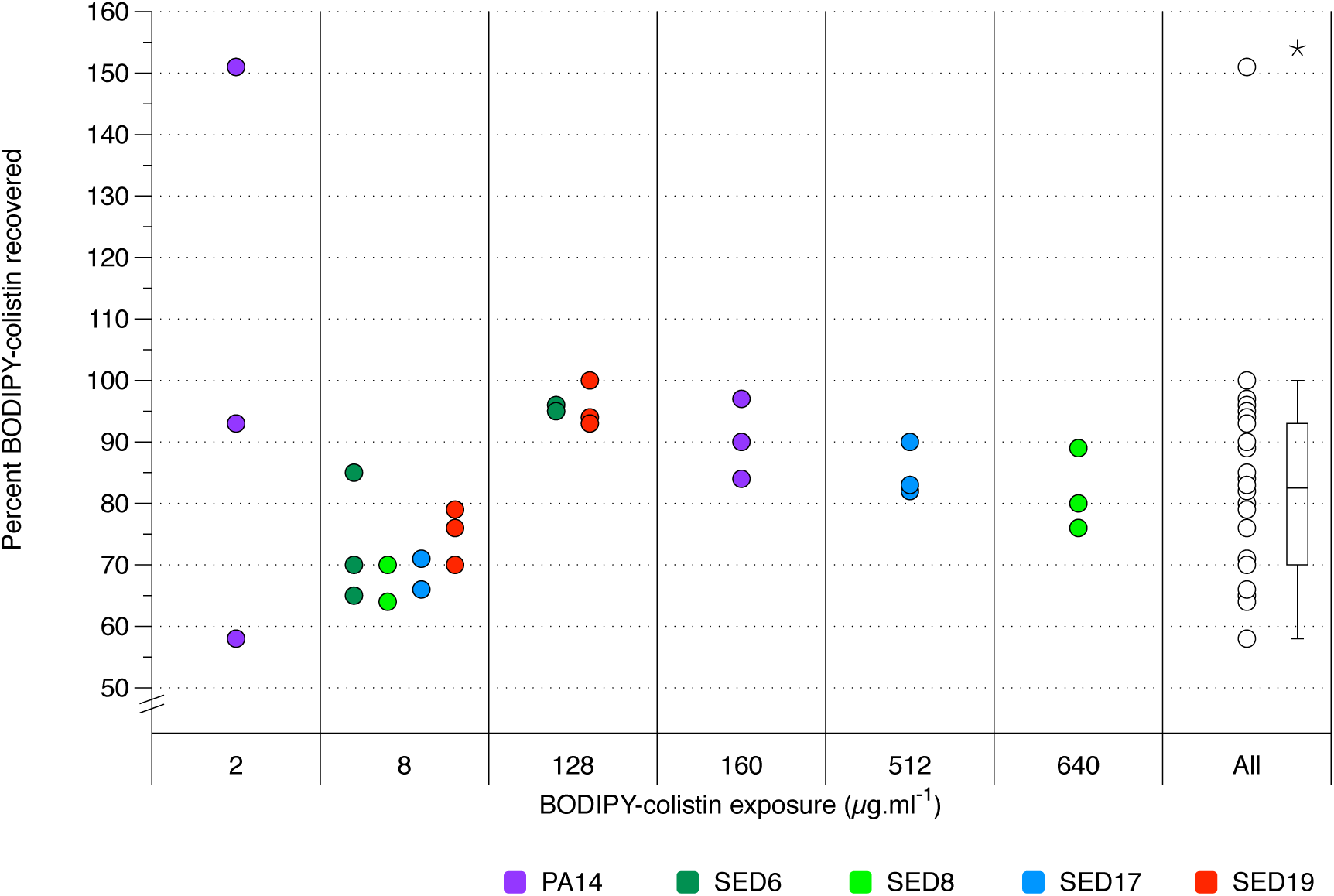
Recovery of initial dose of BODIPY-colistin, as measured by fluorimetry of biofilm homogenate and surrounding SCFM after 18h exposure. Each symbol is one tissue section. Box shows 1^st^ and 3^rd^ quartile with median line, whiskers show interquartile range, asterisk shows outlier. Note that tissues + biofilms were exposed to BODIPY-colistin in a total volume of 1 ml SCFM, therefore concentrations correspond to total µg present. Raw data is supplied in the Data Supplement.

Figure 3a shows the amount of BODIPY-colistin measured in the tissue-associated biofilms for the CF isolates; PA14 is not shown due to the BODIPY quantification becoming unreliable in the 2 µg.ml^-1^ treatment (data are supplied in the Data Supplement). The amount of BODIPY-colistin in biofilms ranged from 12-19% of the total amount recovered from biofilms + surrounding SCFM (means and standard deviations: SED6 13±1%, SED8 16±4%, SED17 15±2%, SED19 13±0%). To calculate the concentration of BODIPY-colistin present in biofilms, the volume of the biofilm was estimated. Conservative estimation of this concentration will be achieved by estimating the largest possible biofilm present on the tissue. Assuming complete coverage of both sides of a 44 mm^2^ tissue section (the average area, see above) by biofilm 100 µm deep (as measured for PA14 in another study using this model, (24)), this suggests a sensible maximum biofilm volume to use is 8.8 µl. Figure 3b shows the predicted BODIPY-colistin concentrations in the biofilms assuming this volume. The concentrations of BODIPY-colistin inside the biofilm following supplementation of the surrounding SCFM at MBC are between 10 and 100X the MBEC measured for the same strain in a Calgary biofilm device using SCFM as the culture medium.

**Figure 3.**
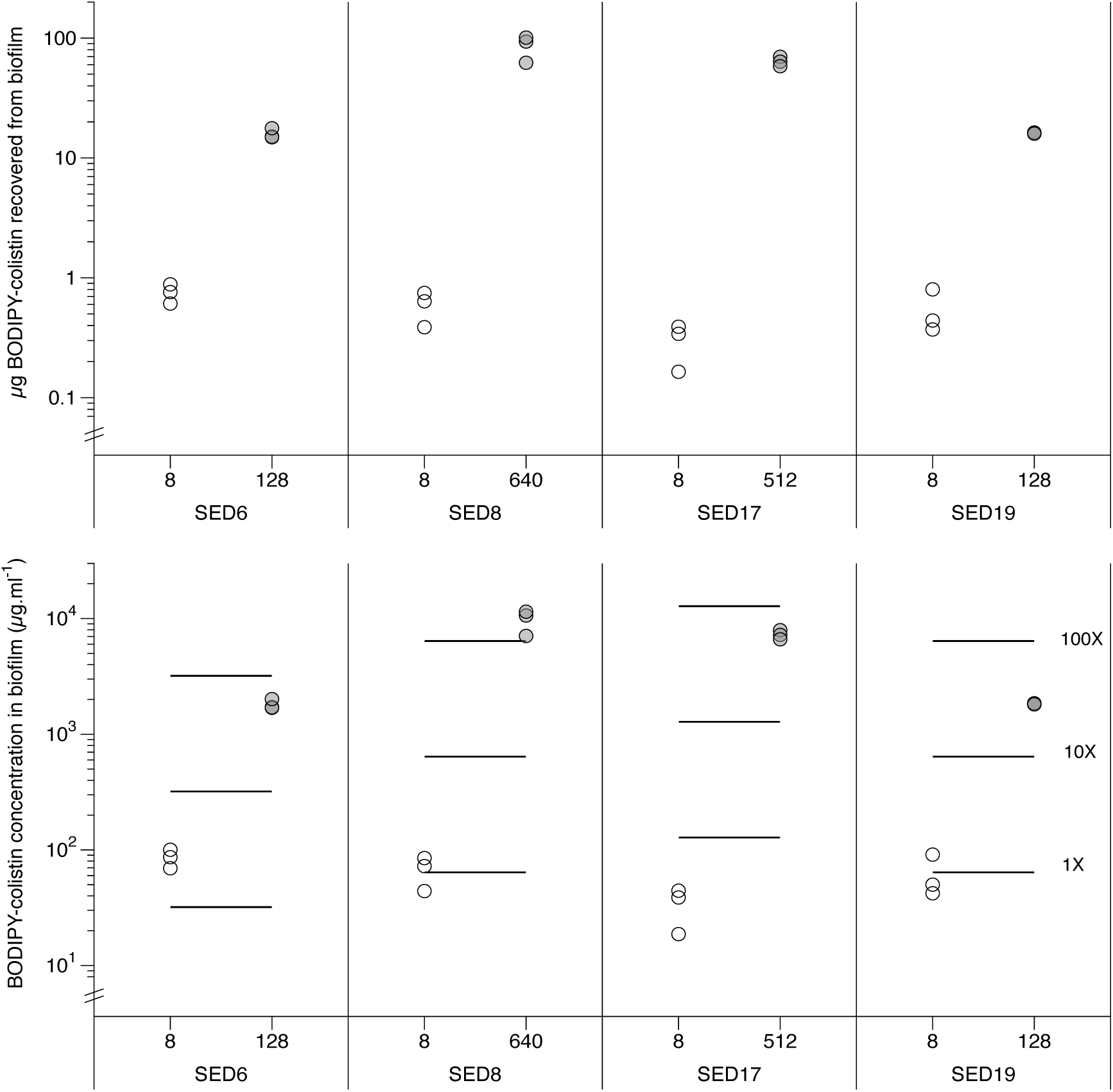
Amount of BODIPY-colistin present in biofilms of CF isolates after 18h exposure, as measured by fluorimetry of biofilm homogenate. Each symbol is one tissue section, numbers on the *x* axis are the sub-inhibitory and MBC exposure doses used. **(a)** µg BODIPY-colistin measured. **(b)** Concentration of BODIPY-colistin in biofilm, assuming a biofilm volume of 8.8 µl. For reference, the lines show 1X, 10X and 100X each strain’s MBEC as measured in the Calgary Biofilm Device using SCFM. Raw data, and data for PA14, are supplied in the Data Supplement.

In-host environments can cue changes in bacterial transcriptome and physiology which will affect sensitivity to antibiotics (e.g. changes in membrane biology, expression of efflux pumps or beta-lactamases (11, 14, 30)). Further, the biofilm mode of growth can itself trigger physiological changes in antibiotic tolerance (e.g. antibiotics that target transcription/translation are not active against quiescent cells deep within biofilms (31)). But a big issue with *in vivo* biofilms is the inability of antibiotics to penetrate the extracellular matrix. Penetration capabilities will depend on the molecular properties of specific drugs, and on the nature of the *in vivo* biofilm matrix. Understanding how the *in vivo* matrix retards drug penetration is a key question in antibiotic choice, dosing and development.

This study shows that colistin, a drug commonly prescribed for *P. aeruginsa* lung infection people with CF, shows very low ability to enter the biofilm matrix when this bacterium is grown as a CF-like bronchiolar biofilm. Combining SCFM with lung tissue to grow CF-like biofilms leads to tolerance of much higher concentrations of colistin than using SCFM with a standard *in vitro* biofilm platform, and this is likely a combination of changes in cellular physiology and the biofilm matrix architecture, as <20% of the labelled colistin to which biofilms were exposed was able to enter the biofilm (these percentages take into account loss of colistin due to adsorption or binding to the lung tissue, glass exposure vial, plastic homogenization tube and/or metal homogenization beads). Peak sputum concentrations of colistin following delivery by inhalation are in the range of 1-40 µg.ml^-1^, declining to 1-10µg.ml^-1^ twelve hours post dose (32, 33), well below the concentrations necessary to observe significant killing of bacteria in entrenched biofilms in our lung model. Colistin dose is limited by its nephrotoxicity. Our results underline the reasons why antibiotics cannot completely clear biofilm infection once it is established: the large doses required to penetrate and kill bacteria in large biofilm aggregates are not physiologically achievable.

EVPL is a tractable and reproducible model for growth of *in vivo*-like *P. aeruginosa* biofilms. Our experiments using EVPL and SCFM alongside standard antibiotic susceptibility testing platforms demonstrate three important microbiological results. First, we confirm that in-host diversity may be important for overall infection AMR: different *P. aeruginosa* clones taken from a single CF sputum sample have different MICs and MBECs *in vitro* in Mueller-Hinton broth and SCFM, and different MBCs in the EVPL biofilm model. Second, growth as *in vivo*-like biofilm in EVPL increases colistin tolerance well beyond what is observed in the Calgary device, even when SCFM is used in place of Mueller-Hinton broth. This underlines the limitations of *in vitro* models of *in vivo* biofilms. Finally, the results confirm that the *in vivo* biofilm prevents free diffusion of colistin to bacterial cells; most colistin administered remains in the SCFM surrounding the biofilm. Clinicians currently have little information beyond planktonic or agar plate MIC to support their choice or dose of antibiotic, so if the EVPL biofilm methodology could be used with microbiological samples from people with biofilm infections (e.g. expectorated sputum) it could facilitate personalised diagnostic AST with greater predictive power, and improved antibiotic stewardship.

These results also suggest that EVPL can be combined with BODIPY-labelling of in-use or novel antibiofilm agents to produce a cheap and simple method for assessing how well these molecules enter the biofilm matrix. Methods such as confocal or super-resolution microscopy, or mass-spectrometry based imaging are already used to visualise how drugs penetrate biofilms, but these require specialised and expensive equipment: this much simpler methodology could speed up biofilm efficacy screening for new antibiotics, and for adjuvants proposed enhance biofilm entry of antibiotics. The main limitation of this approach is unreliable recovery and/or quantitation of BODIPY-colistin at the lowest exposure dose used (2µg.ml^-1^). Further optimisation of the protocol may allow use for lower concentrations of labelled molecules, or alternative approaches such as HPLC or ELISA may need to used when working with low concentrations. The present research focusses on CF, but the combination of an *ex vivo* model using host-mimicking surfaces/media plus fluorescently tagged antibiotics could also be applied to work on other hard-to-treat biofilm infections e.g. ventilator-associated pneumonia (34), chronic obstructive pulmonary disease (35), chronic wounds (36) or medical device infections (37).

Understanding why biofilms are so refractory to treatment and finding new anti-biofilm therapies are priorities in bacteriological research and in industrial R&D (38). Our results underline the extent and diversity of biofilm matrix resistance to antibiotic entry in CF-like biofilm, and provide a platform for more mechanistic exploration of the properties of the *in vivo* biofilm matrix.

The data supplement will be made available with the published version of the manuscript.

## Author statements

FH: Conceptualisation, Methodology, Validation, Formal Analysis, Investigation, Writing – Original Draft Preparation, Project Administration, Funding. ES: Methodology, Validation, Investigation, Writing – Review & Editing. AS: Resources, Writing – Review & Editing. AME: Resources, Writing – Review & Editing.

## Acknowledgements

We thank Akshay Sabnis & Andrew Edwards for BODIPY-colistin; Sophie Darch and Steve Diggle for CF isolates; Niamh Harrington for technical assistance in the lab; John Moat and Anita Catherwood from Warwick Antimicrobial Screening Facility for performing the MIC and MBEC testing; Samantha Westgate & Stefania Fabbri (Perfectus Biomed Ltd). and the National Biofilms Innovation Centre for stimulating conversations which led to this project; and Sheyda Azimi, Steve Diggle, Andrew Edwards, Ramón Garcia Maset & Akshay Sabnis for helpful comments on an early draft of this manuscript. We would also like to acknowledge the help of the Media Preparation Facility in the School of Life Sciences at the University of Warwick, with special thanks to Cerith Harries and Caroline Stewart. This work was supported by an MRC New Investigator Research Grant [grant number MR/R001898/1] awarded to FH. AS is supported by a PhD studentship funded by a Medical Research Council Doctoral Training Award to Imperial College London (MR/N014103/1).

